# Mapping the Cerebello-Hippocampal Circuit: Normative Patterns and Sex-Dependent Connectivity

**DOI:** 10.1101/2025.09.20.677518

**Authors:** Tracey H. Hicks, Jessica A. Bernard

## Abstract

Cerebello-hippocampal (CB-HP) interactions have been implicated in spatial abilities and reinforcement learning, yet their relationship to behavior and differences in connectivity with sex in early adulthood are unclear. Resting CB-HP network patterns have yet to be established in young adults. Mapping the normative resting-state CB-HP connectivity pattern is essential for identifying when CB-HP circuitry becomes behaviorally relevant across the lifespan and for detecting early circuit-level deviations that may precede neurodegenerative or endocrine disruption. We combined resting-state functional MRI (rsfMRI) with measures of cognition in 1,081 healthy young adults (22–37 years, 54% women) from the Human Connectome Project S1200 to map CB-HP functional connectivity (FC), examine sex differences, and define its relationship with performance across episodic memory, visuospatial, and executive function tasks. Region of interest (ROI)-to-ROI FC between the right cerebellum and left hippocampus were quantified. In our CB-HP mapping, we found: 1) negative relationships between the entire HP axis and cerebellar regions: vermis VII; lobules VI, VIIb, and VIII; Crus I & II, and 2) positive relationships between most of the HP axis and ventral/medial cerebellar regions. We revealed sex differences in CB-HP such that females had greater FC than males between anterior to mid-hippocampal regions and medial cerebellar regions: vermis IV-VI; lobules VI, VIII, IX; and Crus II. We predicted CB-HP FC patterns would show strong positive associations with cognitive measures (i.e., episodic memory, visuospatial processing, working memory, and executive function); however, we did not find any associations after multiple comparisons correction (pFDR > 0.05). Together, our findings detail a functional atlas of the CB-HP circuit in young adulthood and highlight sex differences within. Our results provide a foundation for understanding functionally-based gradients between these two regions.

## Introduction

Converging animal and human evidence shows that cerebellar (CB) contributions extend well beyond motor abilities and encompass a wide variety of cognitive measures including spatial abilities and aspects of memory^1–4^; tasks which have classically been attributed to the hippocampus (HP). On a cellular level, tract-tracing in mice revealed bidirectional connectivity along CB–HP pathways that displayed synchrony during tasks requiring spatial navigation and motor control, providing direct evidence for the role of CB-HP in goal-directed behavior^5^. Further, optogenetic experiments demonstrate a causal cerebellar influence on hippocampal dynamics^6,7^. Specifically, cerebellar photo-stimulation suppresses spontaneous HP seizures in mice with temporal lobe epilepsy^6^ and acutely reshapes hippocampal CA1 oscillations and place-cell coding^7^. Combined, these findings provide compelling evidence for CB-HP pathways in the rodent brain, with impacts on cognitive performance.

Functional MRI (fMRI) in humans has expanded our understanding of the CB-HP circuit in the context of cognition. Co-activations between CB and HP were demonstrated when participants predict the future position and timing of moving targets^8^, decode egocentric versus allocentric changes in visual scenes^9^, and memorize action sequences during virtual-maze navigation^10^. Most recently, analyses that allow a high-dimensional matrix of cerebellar-cortical connections to self-organize into a few smooth axes (manifold-learning analyses) showed that stronger CB-HP (and broader CB-cortical) integration accelerates reward-based motor reinforcement learning in young adults^11^. Notably, these studies probe isolated tasks and limited region sets, leaving the overall topology of the CB–HP circuit largely undescribed. To date, no study has produced a normative, regionally resolved resting-state map of CB-HP connectivity in healthy young adults. Available rsfMRI work has focused on middle-aged and older cohorts^12^ or examined the circuit only indirectly^13^. This gap constrains translation, because without a baseline FC blueprint, it is difficult to detect early circuit-level deviations tied to hormonal transitions or emerging neuropathology.

The functional topology of the hippocampus is organized along smooth functional gradients rather than discrete anatomical regions^14–16^. The anterior hippocampal has been strongly associated with memory and semantic processing^14,15,17^; whereas, the posterior hippocampus has been linked to spatial navigation^15^. Yet across these demonstrations investigational methods are varied (e.g., manifold-learning, meta-analysis, task-based fMRI), leaving the baseline topology of hippocampal (and CB-HP) connectivity in neurotypical young adults largely uncharted. Baseline gradient maps in neurotypical young adults provide the coordinate system against which developmental shifts, sex differences, and disease-related changes can be quantified; they anchor hippocampal subregion definitions, improve cross-study comparability, and enhance sensitivity when testing for subtle changes in later life.

Sex differences present another underexplored facet of the CB-HP circuit. Recently, sex differences were found in functional brain organization primarily in the default mode network, striatum, and limbic network^18^. Interestingly, machine-learning work shows that connections within visual, attentional, and temporoparietal networks predicted both crystallized and fluid cognitive abilities in young males and females alike, with only modest sex-specific nuances^19^. Variation in brain function by sex stands to reason as sex steroid hormones differ by biological sex and modulate brain function^20–22^.

Notably, there is a higher density of estradiol and progesterone receptors in the cerebellum and hippocampus^23^. Current data suggest that cognitively normal middle-aged and older adults have not demonstrated sex differences in CB-HP FC^12,13^. However, this has not been directly investigated in young adults. Clarifying whether CB-HP FC and its cognitive correlates are equivalent in neurotypical young adult females and males will refine normative baselines against which lifespan or clinical deviations can be benchmarked.

The present study aims to provide a high-resolution, regionally specific map of resting-state functional magnetic resonance (rsfMRI) connectivity between the cerebellum and hippocampus in healthy young adults, establishing a normative reference for later lifespan and prodromal neurodegenerative comparisons. Available evidence for this circuit in humans in largely derived from visuospatial task paradigms^8–10^ rather than systematic rsfMRI characterization. A regionally resolved approach is essential because both structures exhibit pronounced functional heterogeneity. Specifically, meta-analytic functional connectivity (FC) evidence has shown a graded anterior–posterior specialization of the hippocampus, with anterior portions more strongly associated with semantic and associative memory and posterior hippocampus with spatial abilities amongst several cognitive measures(i.e., episodic encoding and retrieval, semantic retrieval, working memory, spatial navigation, simulation/scene construction, transitive inference, and social cognition tasks)^14^. Similarly, the cerebellum shows a gradient of region specific participation in a wide variety of cognitive operations (e.g., autobiographical recall, working memory, executive function^1,3,24^. Establishing whether hippocampal subregions couple preferentially with specific posterior cerebellar territories in early adulthood is critical for interpreting later alterations attributed to hormonal change, cognitive aging, or Alzheimer’s disease–related biomarker status^25^. By integrating anatomically informed regions of the hippocampus and cerebellum, this work seeks to delineate the foundational topology of the “hippobellum” network before compensatory or pathological processes emerge.

Here, we used the Human Connectome Project (HCP) S1200 to close this gap. We examined over 1,000 young adult participants with resting-state fMRI data to map the CB-HP circuit in young adults, examine sex differences in CB-HP FC, and assess cognitive associations with CB-HP connectivity. Recent work has shown functional connectivity (FC) between CB and HP^8–11,13^; however, the CB-HP circuit and its associations with region specific cognitive performance has yet to be examined at this level of depth (several ROIs covering each anatomical region) nor with such a large sample of young adults. Without a characterization of CB-HP FC, early circuit-level shifts linked to hormonal change or insidious neuropathology are hard to spot, limiting translational progress. Further, determining whether stronger resting CB-HP FC predicts performance on visuospatial, episodic-memory, inhibitory-control and working-memory tasks would advance models of cognition.

Leveraging data from 1,081 HCP participants, the present study first sought to map in detail the connectivity between the cerebellum and hippocampus, taking a detailed regional approach to account for functional heterogeneity and anatomical variance in both regions. We also test the following hypotheses: 1) that there are sex differences in CB-HP FC; 2) that stronger CB-HP FC is associated with better performance on measures of visuospatial processing, episodic-memory, working memory, and executive function; and 2) CB-HP FC and its cognitive associations would differ significantly between young-adult females and males.

## Method

Our study used the HCP S1200 fMRI dataset, which was comprised of 1,113 young, healthy adult subjects (age 22-37) with MRI data. Details of the dataset can be found in the HCP Data Release Reference Manual^26^ and the design of the experiment is summarized by Van Essen et al.^27^.

### Study sample

Participants (*n* = 1,081) from the HCP were included in our analyses. Participants were included if they had two rs-fMRI nifti files and one T1 file from the first scan session. 32 participants were excluded for incomplete files. Our sample was 54% female (n=585 F) with an age range of 22-37 years (*M* = 28.78 years, SD = 3.70), and an education level with a range of 11-17 years (*M* = 14.90 years, SD = 1.90). Most participants completed every cognitive task examined in this study; however, we included a breakdown of participant numbers by task (**Table 1**). Each task was evaluated separately with the maximum number of participants possible in each analysis (i.e., using every participant with available data). All cognitive tasks used education level as a control in our analyses and 2 participants were excluded for missing education level data. Additional recruitment and exclusion criteria for the HCP are reported here (humanconnectome.org/storage/app/media/documentation/s1200/HCP_S1200_Release_Reference_Manual.pdf#page = 125.19). Handedness of participants in our sample is as follows: 86% were right-handed, 9% were ambidextrous, and 6% were left-handed per self-report on the Edinburgh Handedness Inventory^28^. While demographics such as ethnicity, race, and genetic/environmental sibling data were collected in HCP, those data were categorized as restricted data elements. These restricted demographics will not be reported in this study due to their sensitive nature. Further, race and ethnicity were not considered in this study as information about socio-economic status is better indicated for causal inferences in neuroimaging^29^. Notably, genetic and environmental kinship was accounted for statistically (explained further in *Statistical Analyses*).

**Table 1.** Number of participants are listed for the full sample and each task after removing participants without education level data.

| Participants per Task |  |
| --- | --- |
| Full sample | 1081 |
| Short Penn Line Orientation | 1073 |
| Picture sequence memory | 1078 |
| Penn Word Memory Test | 1073 |
| Flanker task | 1079 |
| List Sorting Working Memory | 1079 |

### Cognitive Measures

Participants were administered computerized cognitive tests from two standardized batteries, the NIH Toolbox (version 1.11)^30,31^, and the University of Pennsylvania Computerized Neuropsychological Test Battery^32^. A priori cognitive measures were selected to evaluate CB-HP FC associations with these tasks.

#### Visuospatial processing

The Variable Short Penn Line Orientation Test^32^ assesses visuospatial processing^26^. This task presents participants with two lines on each trial: a fixed red reference line and a blue line that the participant can rotate. Using keyboard keys to turn the blue line clockwise or counterclockwise, the participant tries to align it so that the two lines are parallel. Although the center-to-center distance between the lines remains constant, their on-screen positions change from trial to trial, and the blue line appears in either a short or long version, whereas the red line’s length never varies. The test comprises 24 alignment trials. As several studies have demonstrated links between visuospatial performance and CB-HP pathways^8–11^, performance on this measure was expected to be positively associated with CB-HP FC.

#### Episodic Memory

The Penn Word Memory Test^32^ (Form A) evaluates verbal episodic memory^26^. Participants are shown 20 words and asked to remember them for a later memory test. Later, they are shown 40 words, 20 target and 20 new, memory-matched foils. They are tasked at judging each word’s prior occurrence by selecting one of four confidence options: “definitely yes,” “probably yes,” “probably no,” or “definitely no.”

Non-verbal episodic memory is assessed using the NIH Toolbox Picture Sequence Memory Test^30^. This measure includes the acquisition, storage and effortful recall of new visual information. It involves recalling an increasingly lengthy series of illustrated objects and activities that are shown on a computer screen in a specific order. Both measures were used to test the hypothesis that CB-HP FC is associated with episodic memory function.

#### Working Memory

The List Sorting Working Memory Test from the NIH Toolbox assesses working memory^30^. This task involves both information processing and storage. Participants sequence several visually and orally presented stimuli (foods and animals) into size order. Higher scores on each of these indicate higher levels of working memory. Performance on this task is a compelling correlate for CB-HP FC because it requires people to hold a short series of items online and then reorder that series. This cognitive process simultaneously taps the hippocampus’ capacity to tag each item with its temporal-context position— crucial for remembering the original order^33^, and the cerebellum’s inner-speech and sequencing mechanisms that refresh and manipulate the phonological list^34^.

#### Executive Function

The NIH Toolbox Flanker Inhibitory Control and Attention Test measures attention and inhibitory control^30^. Participants are presented with five arrows on each trial and must indicate, via left- or right-key presses (or touchscreen buttons), the direction of the central arrow while ignoring the flanking arrows. After a brief practice block with feedback, adults complete 40 test trials (20 congruent, 20 incongruent) in pseudorandom order: a fixation cross appears for 400 ms, the arrow array remains on-screen for up to 1,600 ms or until response, and a 350 ms blank interval separates trials. Performance is summarized as an accuracy-adjusted reaction-time score (mean = 100, SD = 15), providing a standardized index of inhibitory control and attention. We included this measure as there is evidence that resolving distractor interference engages cerebellar circuits that support executive attention^21^ and suspect hippocampal mechanisms may also be engaged in this process^36^.

### Imaging acquisition and Processing

In this study, rsfMRI data from the Human Connectome Project (HCP S1200 release) were used^27^. Two, 15-minute runs of rsfMRI data were acquired in the first session on a 3 T Siemens Connectome-Skyra scanner with gradients customized for the HCP. Briefly, a gradient-echo echo-planar imaging (EPI) sequence (1,200 time frames; TR = 720 ms; TE = 33.1 ms; flip angle, 52°; 2.0 mm isotropic voxels; multiband factor, 8) was used in each run of rsfMRI data. Data was collected with the participants’ eyes open. Preprocessed resting-state T1w structural and resting-state functional MRI (rsfMRI) data from the S1200 Release of Human Connectome Project (WU-UMN HCP Consortium) were used for analysis^26^.

All rsfMRI data were processed with the HCP minimal preprocessing pipelines (v3), which use FSL 5.0.6, FreeSurfer 5.3.0-HCP, and Connectome Workbench v1.1.1 and comprise structural (PreFreeSurfer, FreeSurfer, PostFreeSurfer) and functional (fMRIVolume, fMRISurface) pipelines. The fMRIVolume pipeline corrects gradient distortions, performs motion correction with FLIRT using the single-band reference (SBRef) volume as target, applies TOPUP-based EPI distortion correction using spin-echo field maps, registers the EPI to the T1w image with a customized FLIRT BBR + bbregister refinement, resamples the data to MNI space in one spline interpolation step that combines all transforms, and applies global mean intensity normalization (mean = 10,000) and bias-field reduction, avoiding any additional spatial smoothing. The subsequent fMRISurface pipeline maps the preprocessed 4D volume to the CIFTI grayordinates space (91,282 grayordinates; 32k fs_LR surface per hemisphere + subcortical voxels) via cortical ribbon-based volume-to-surface mapping, applies surface (2 mm FWHM) and parcel-constrained subcortical smoothing, and outputs a dense timeseries suitable for connectivity analyses. For additional details, please refer to comprehensive HCP preprocessing workflows^37^ (humanconnectome.org/storage/app/media/documentation/s1200/HCP_S1200_Release_Reference_Manual.pdf#page = 125.19).

The remaining section covering imaging analyses use standardized text to ensure consistency and reproducibility in the research protocol. These standardized descriptions can be found in other publications from our laboratory^12,38,39^ and align with current best practices in the field, providing a clear and detailed framework for the procedures undertaken.

Statistical analyses were conducted using the CONN toolbox (22v2407)^40^. This involved additional processing to eliminate noise and artifacts and enhance data quality. Denoising in CONN consists of several stages, such as removing motion signals and regressing out confounding signals (e.g., signals from ventricles, white matter, and global signals). A 0.008-0.099 Hz (default) bandpass filter was applied to eliminate high-frequency noise. The denoising step is crucial for enhancing the quality of FC data by minimizing artifacts and enhancing the ability to detect genuine FC patterns in the brain.

Resting-state FC analyses focused on regions of interest (ROIs) in both the hippocampus and the cerebellum (**Figure 1**). The hippocampal seed regions were loosely based on coordinates derived from Grady’s meta-analysis on hippocampal function^14^. These coordinates helped determine anatomical hippocampal boundaries and the coordinates were shifted from the original values to accommodate a new center for 5mm spherical seeds in MNI space. That is, certain coordinates from Grady^14^ fell on the outer edges of the hippocampus and to cover 3-dimensional space within the boundaries of the hippocampus, we had to shift certain coordinates to account for a 5mm spherical seed. Cerebellar seeds were determined by mapping dorsal, ventral, rostral, caudal, medial, and lateral boundaries of the right cerebellum in MNI space. Please refer to **Table 2** for coordinates for all our seeds. Following this step, we used the oro.nifti and spatstat packages in R Studio to create a non-overlapping grid in 3 dimensional space of 5mm spherical seeds covering the right cerebellum^41,42^. As our aim for this study was to cover the entirety of the right cerebellum and left hippocampus, seeds were created to cover the 3-dimensional anatomical regions in MNI space.

**Figure 1.**
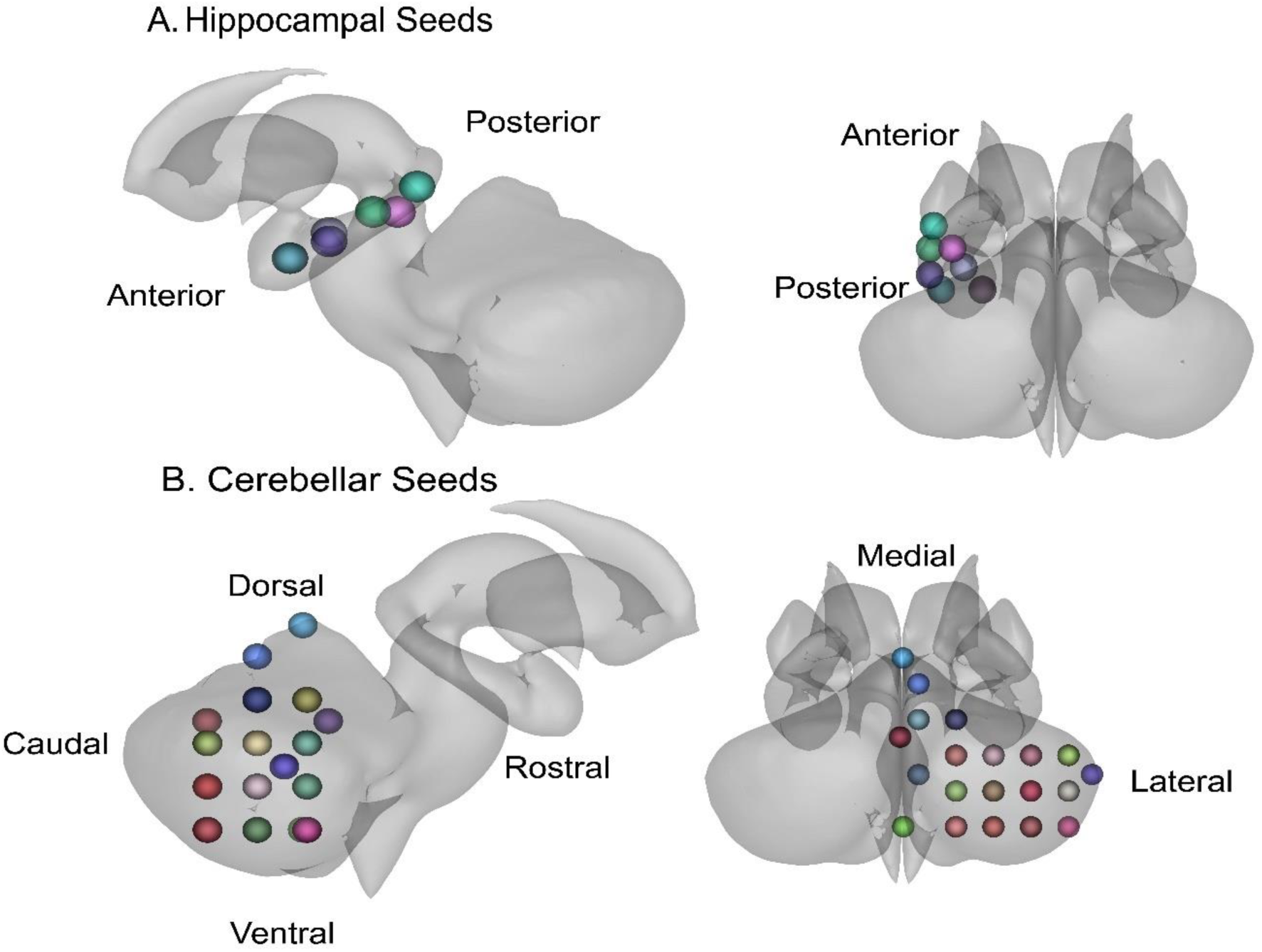
These figures illustrate several regions of interest (ROIs) used in this study. Seed colors are helpful for differentiating seeds, but do not hold additional meaning. **A**. Hippocampal seeds are displayed with directional indicators of the anterior and posterior long-axis. **B**. Cerebellar seeds are displayed with directional indicators for the right cerebellar hemisphere. The caudal region of the cerebellum is better visualized in a more detailed representation of the ROI seeds in **Supplementary Figure 1**.

**Table 2.**
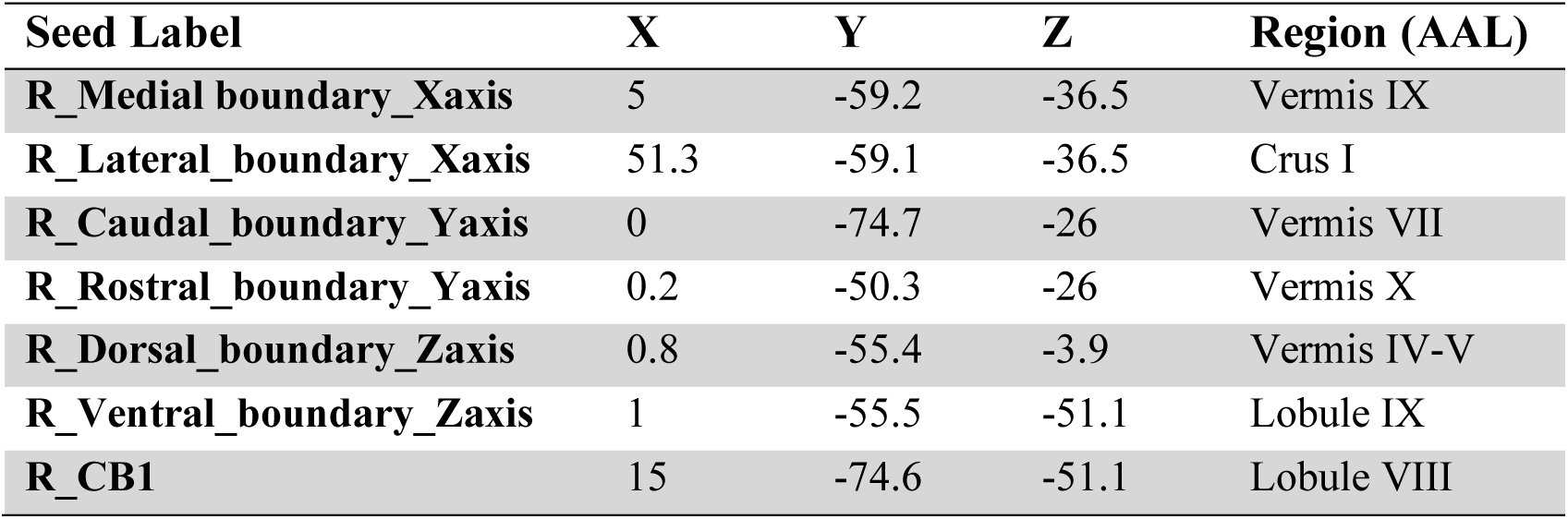
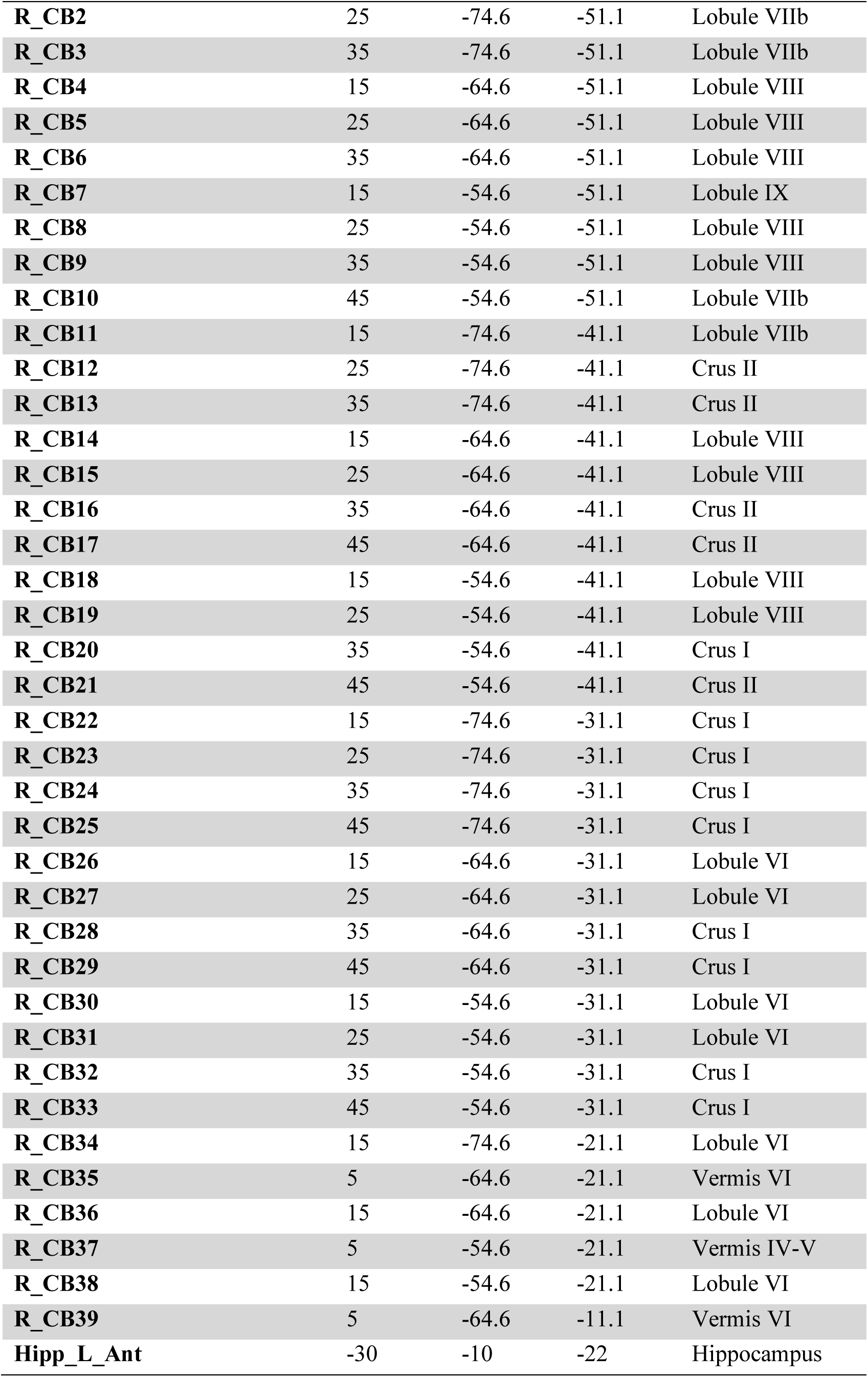
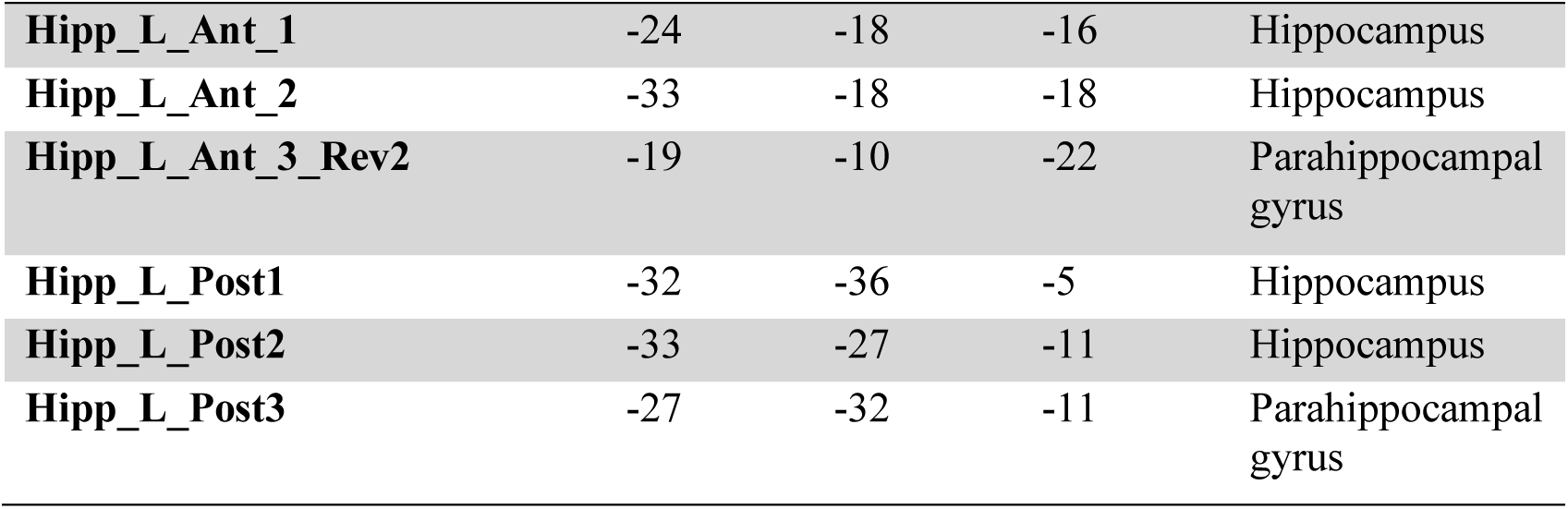
Cerebellar and Hippocampal Coordinates for 5mm Spherical Seeds.

The investigation was limited to the right cerebellar hemisphere and left hippocampus, to mitigate multiple comparisons and ensure examination of cross lateralized connections. Additionally, these cross-lateral regions were chosen as right cerebellar to hippocampal FC relationships have been shown in the context of aging adults (middle-aged to older adults)^4,43^. Lastly, the left hippocampus and right cerebellum may show early indications of neuropathology in aging. In patients with mild cognitive impairment (MCI) and Alzheimer’s disease (AD), there is a consistent pattern of greater left-than-right hippocampal atrophy, with great asymmetry in MCI than AD, indicating the left hippocampus as more vulnerable to neurodegeneration as compared to the right^44^. The right cerebellum displays lower connectivity with cortical regions in amnestic mild cognitive impairment patients as compared to healthy controls^45^. While this study investigated young healthy adults, our overarching intent is to apply the baseline mapping here to identify preclinical neuropathology in aging adults or in other neurological illness or psychopathology impacting these regions.

For ease of interpretation, we used the automated anatomical atlas (AAL) in the label4MRI package in R to approximate anatomical regions for each seed^46^ which is specified in **Table 2**. Our seeds are visualized in a simpler, easier to view, rendering in **Figure 1**, and we have created a more in-depth representation in **Figure 2**. These seeds have also been used in a study of the CB-HP circuit in middle-aged and older adults^12^; our intention was to run parallel analyses with the same seeds across large, healthy adult cohorts to characterize the CB-HP circuit across the lifespan.

**Figure 2.**
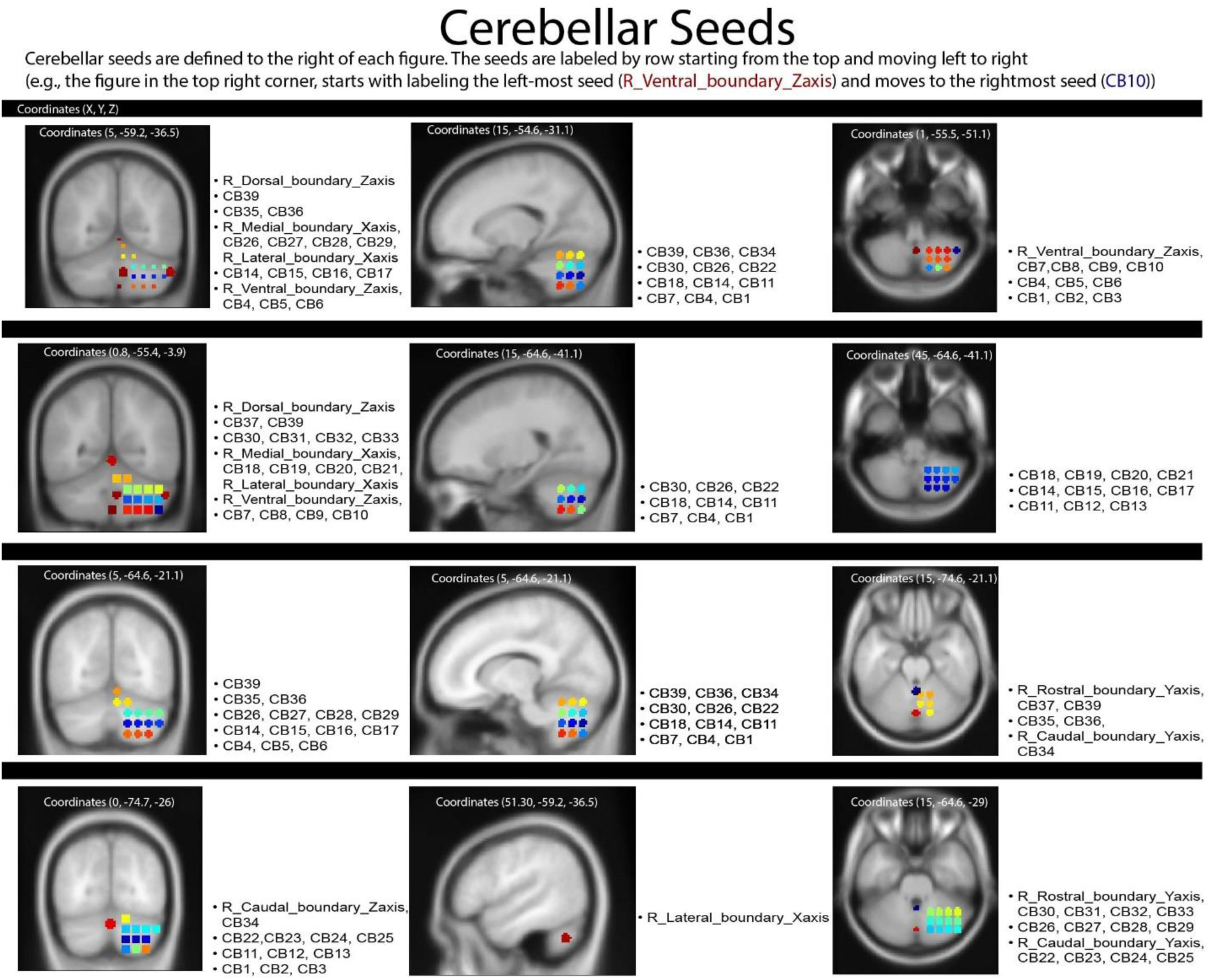
Cerebellar Seeds Rendered in marsbar.

### Imaging and Statistical Analyses

CB-HP FC was examined via ROI-to-ROI correlations using the CONN toolbox. For each subject we extracted all pairwise ROI-to-ROI Fisher-Z transformed bivariate correlation connectivity values into a correlation matrix across 52 ROIs.

A number of participants in HCP S1200 were members of the same family and thus have shared genetic and environmental variance. In our sample, 86% of participants were related; however, additional details on relatedness (e.g., twin zygosity, maternal versus paternal relatedness) will not be divulged due to the restricted nature of these data. We used Permutation Analysis of Linear Models (PALM v. alpha122; https://web.mit.edu/fsl_v5.0.10/fsl/doc/wiki/PALM.html) to permute the data using exchangeability blocks designed to separately capture the unique variance within monozygotic twins, dizygotic twins, and non-twin siblings, while simultaneously accounting for family structure. Exchangeability blocks were generated with the hcp2blocks script provided by the developers of PALM (retrieved from https://github.com/andersonwinkler/HCP). All FC analyses underwent PALM analysis using the aforementioned Fisher z-transformed matrices from CONN Toolbox.

Through PALM, we first investigated and defined the CB-HP circuit in young adults via one-sample t-statistic for each edge (testing mean connectivity ≠ 0). Sex differences in age and education were analyzed with Welch Two Sample t-tests (separately). Then we assessed sex differences in CB-HP FC with a permutation general linear model (GLM) in PALM using a design matrix (1= Female, 0 = Male). GLMs in PALM also evaluated cognitive performance (i.e., Variable Short Penn Line Orientation, Penn Word Memory Test total, Penn Word Memory Test reaction time, Picture sequence memory, Flanker Task, and List Sorting Working Memory) with CB-HP FC (respectively) while controlling for age and education level. Lastly, GLMs in PALM evaluated interactions between sex and cognitive performance with CB-HP FC while controlling for age and education level. Following PALM analysis, R Studio was employed to filter only unique CB-HP connections (314) from the larger full matrix with 2,704 connections which included both within-CB and within-HP connections. We limited our analyses in this way as our goal and hypotheses are focused on connectivity between the CB and HP. False discovery rate (FDR) correction was then applied to correct for the 314 ROI-to-ROI pairs in each analysis at *p* < .05.

For statistical analyses of the age, education level, sex, and behavior results, we used R Studio (v2024.04.2.764). First, demographic (i.e., age and education level) differences by sex were assessed via Welch Two Sample t-tests. We then investigated sex differences in cognitive performance (i.e., Variable Short Penn Line Orientation, Penn Word Memory Test total, Penn Word Memory Test reaction time, Picture sequence memory, Flanker Task, and List Sorting Working Memory) via analyses of covariance (ANCOVAs) which controlled for age and education level. FDR correction was applied to correct for 6 comparisons.

## Results

For the following results, please refer to **Figure 1** as a reference for directionality regarding both the hippocampus and cerebellum.

### Demographics

Our sample was 54% female. Welch Two Sample t-tests revealed significant sex differences in age (t(1047.8) = −7.345; *p* <.0001) in which women were significantly older than men in this sample. Education level was not significantly different by sex (t(1054.9) = −1.602; *p* =.1095). **Table 3** has demographics broken down by sex and denotes variables with significant sex differences. For additional demographic details please refer to the Method section, Study Sample subsection.

**Table 3.**
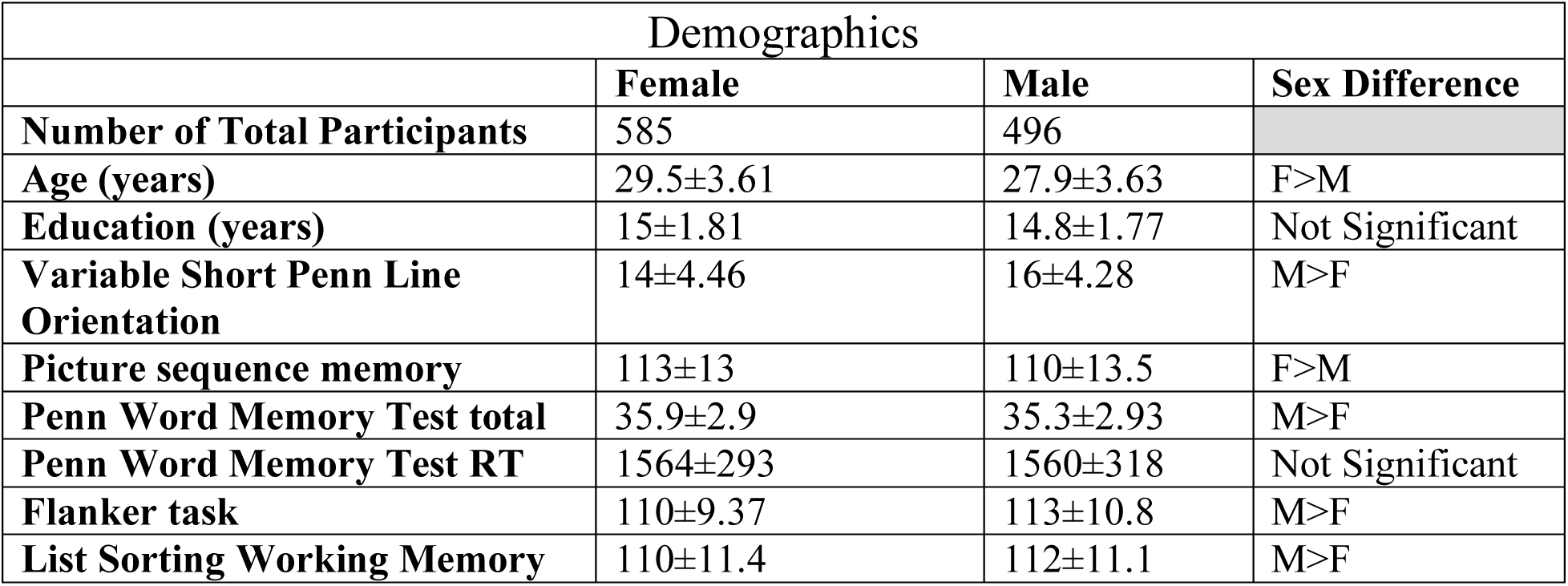
Demographic means and standard deviations each variable, categorized by sex.

### CB-HP Connectivity Mapping

Our mapping of CB-HP FC largely revealed negative relationships between the entire HP axis and cerebellar regions: vermis VII; lobules VI, VIIb, and VIII; Crus I & II (pFDR<.05; **Table 4**, **Figures 3 & 4**). That is, connectivity between most CB-HP pairs was lower than our null hypothesis of zero FC when accounting for kinship. Notably, several CB-HP connections showed positive relationships across most of the HP axis and ventral/medial cerebellar areas (pFDR<.05; **Table 5**, **Figures 3 & 5**). Notably, two posterior HP seeds (Hipp_L_Post1 & Hipp_L_Post2) frequently showed negative or no correlations in the same cerebellar regions. Notably, **Figures 4 & 5** are solely intended for conceptual purposes; they represents data prior to correction for kinship. Only ROI pairs that exhibited strong negative relationships in CB-HP FC after kinship- and FDR-correction are displayed here; however, intra-regional correlations are also visualized due to software limitations in excluding those relationships.

**Figure 3.**
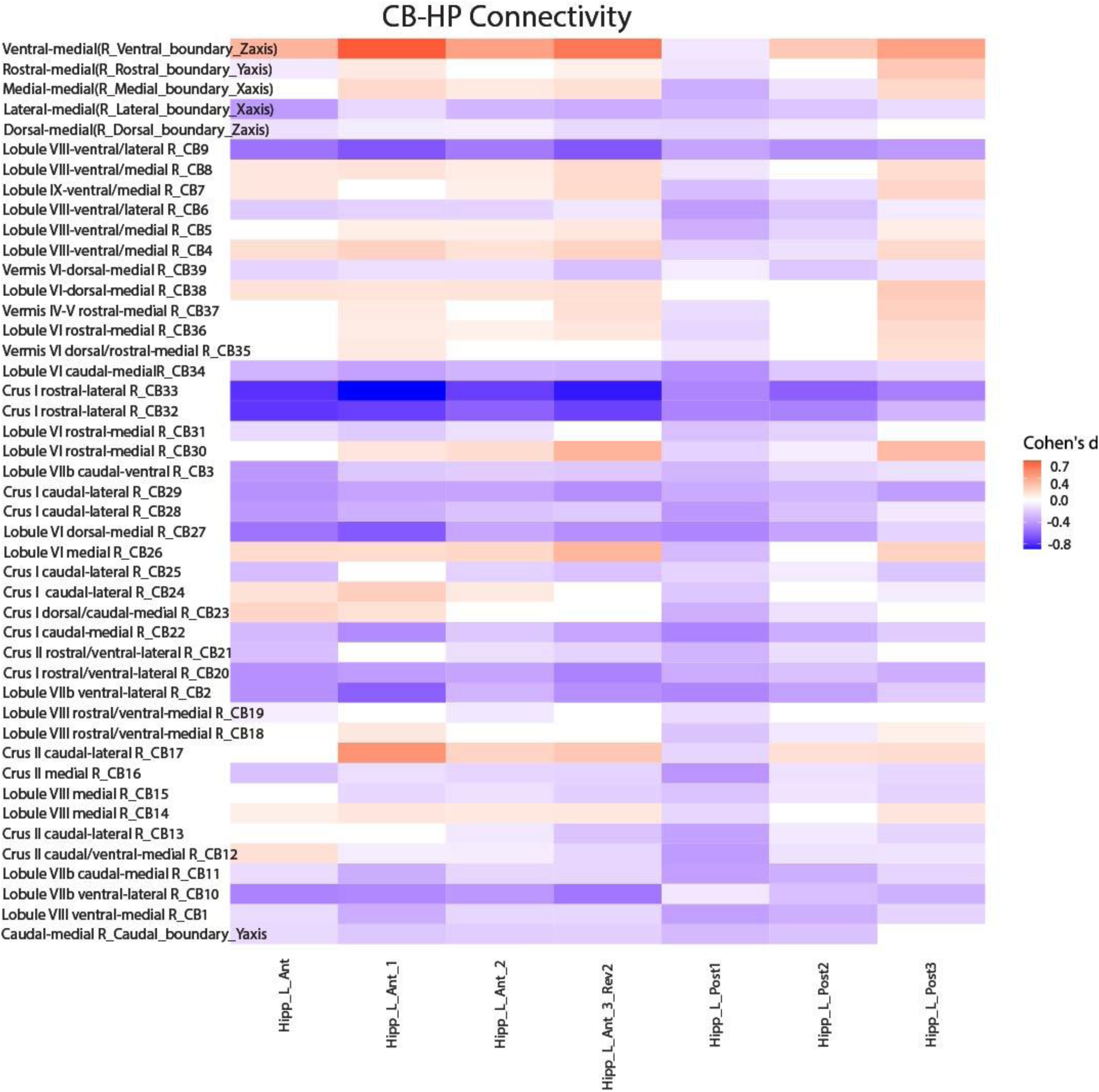
This heatmap displays the region of interest (ROI) pairs used in this study. Red indicates positive connectivity between the two regions and blue indicates negative connectivity. Values were plotted using Cohen’s d to illustrate the effect size for each connection while maintaining directionality.

**Figure 4.**
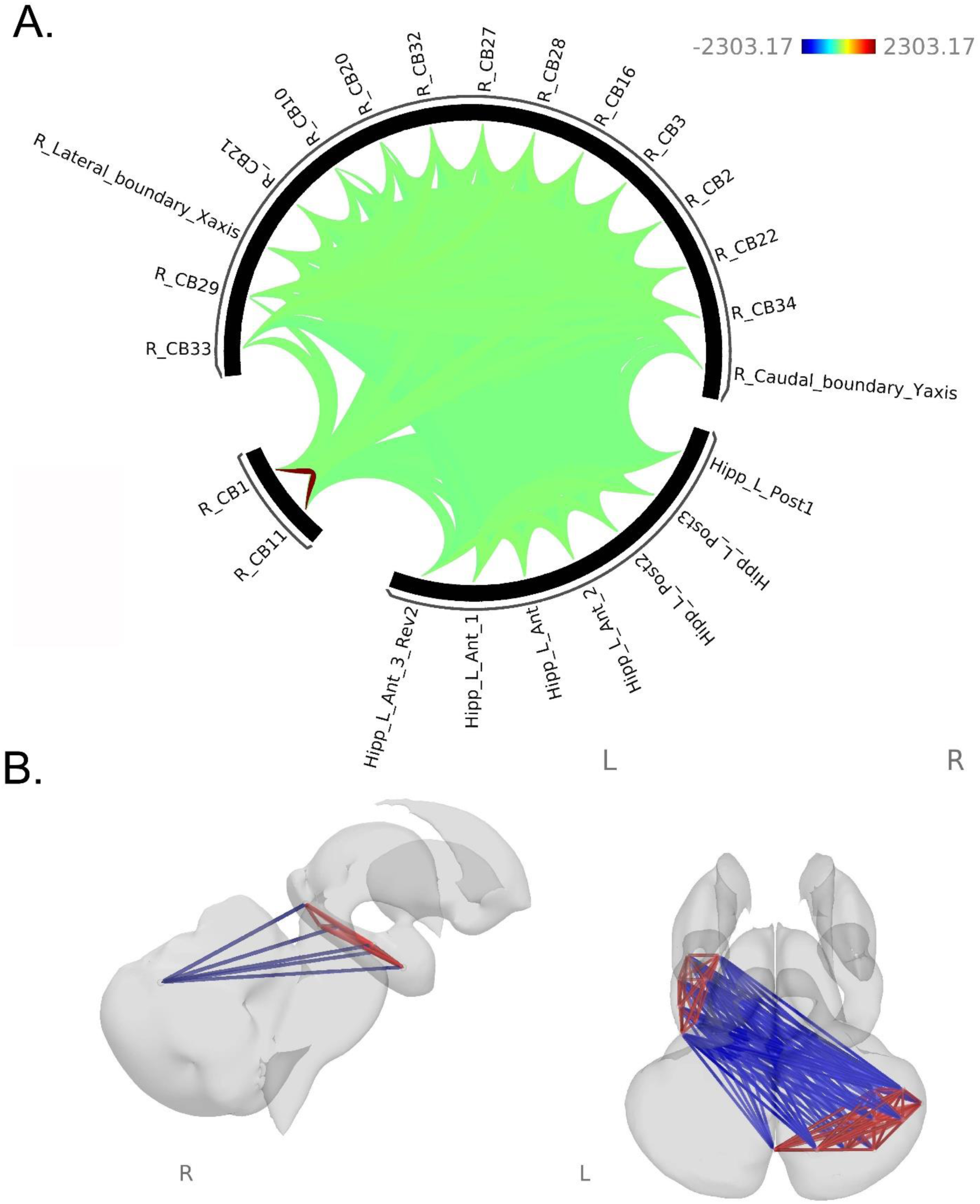
Visualization of negative patterns of CB-HP functional connectivity (FC) across participants. **A**. ROIs are shown in an FC ring where green displays lower FC than the null hypothesis of zero and red is greater FC than zero. **B**. ROIs are shown on a subcortical model where blue displays negative FC relationships and red shows positive relationships. This figure is solely intended for conceptual purposes; it represents data prior to correction for kinship. Only ROI pairs that exhibited strong negative relationships in CB-HP FC after kinship- and FDR-correction are displayed here; however, intra-regional correlations are also visualized due to software limitations in excluding those relationships.

**Figure 5.**
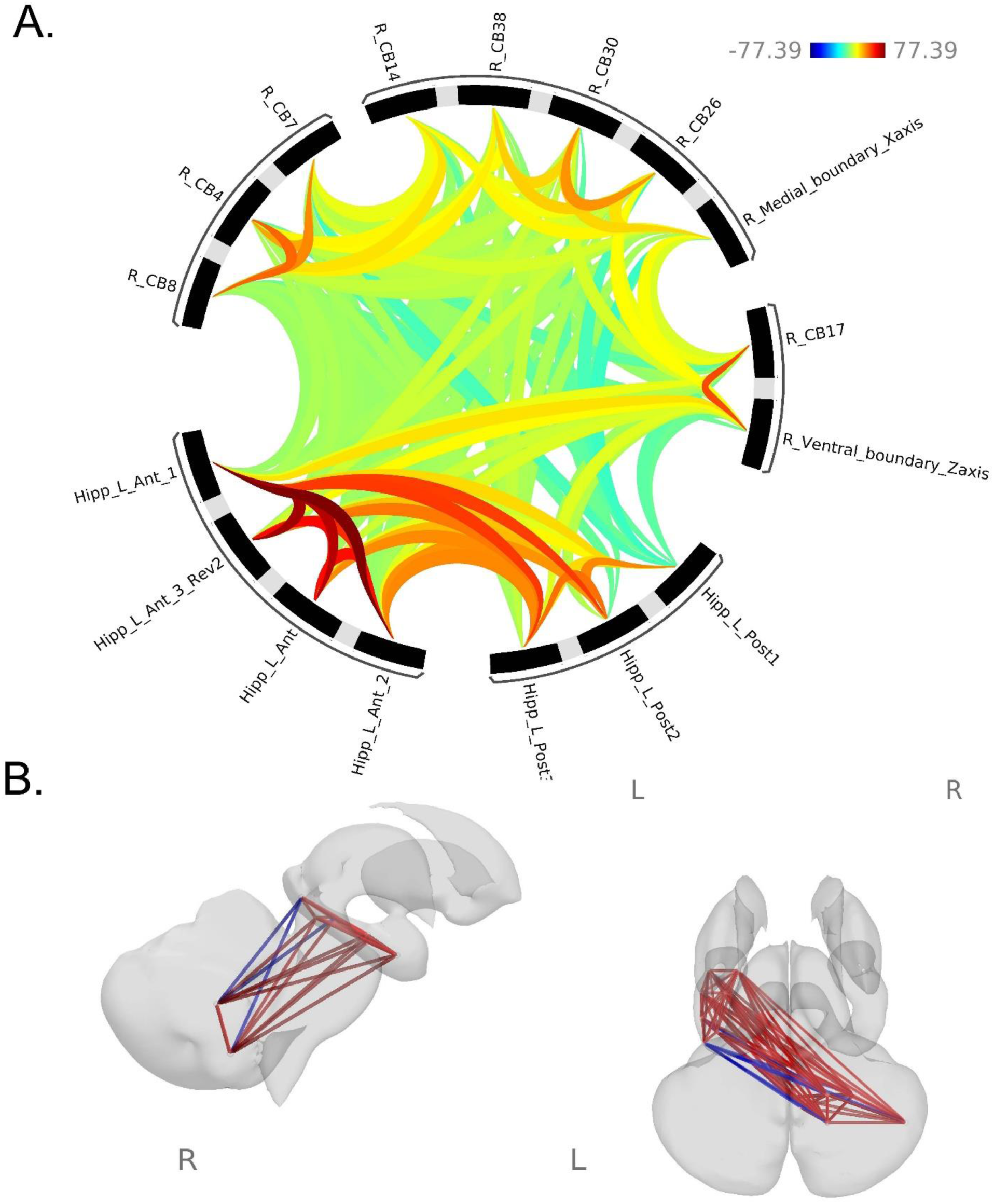
Visualization of negative patterns of CB-HP functional connectivity (FC) across participants. **A**. ROIs are shown in an FC ring where orange-red displays greater FC and blue displays lower FC than our null hypothesis of zero. **B**. ROIs are shown on a subcortical model red shows positive FC relationships and blue displays negative relationships. This figure is solely intended for conceptual purposes; it represents data prior to correction for kinship. ROI pairs that exhibited strong positive and one strong negative (most posterior hippocampal seed) relationships in CB-HP FC after kinship- and FDR-correction are displayed here; however, intra-regional correlations are also visualized due to software limitations in excluding those relationships.

**Figure 6.**
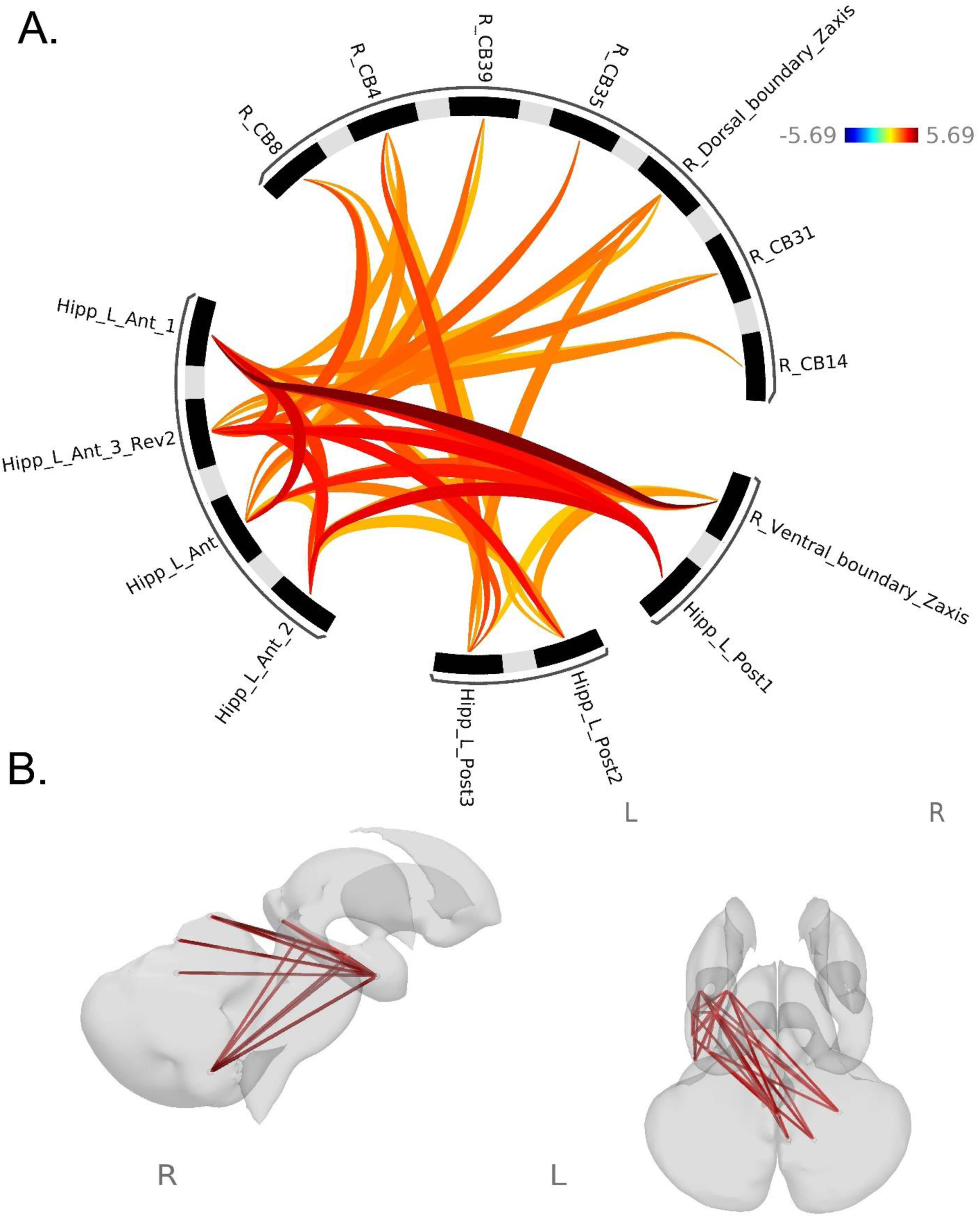
Visualization of sex differences in CB-HP functional connectivity (FC). **A**. ROIs are shown in an FC ring where orange-red displays greater FC in females as compared to males. **B**. ROIs are shown on a subcortical model red shows greater CB-HP FC in females as compared to males. This figure is solely for conceptual visualization purposes; it represents data prior to correction for kinship. Only ROI pairs that exhibited significant relationships in CB-HP FC after kinship- and FDR-correction are displayed here; however, intra-regional correlations are also visualized due to software limitations in excluding those relationships.

**Table 4.**
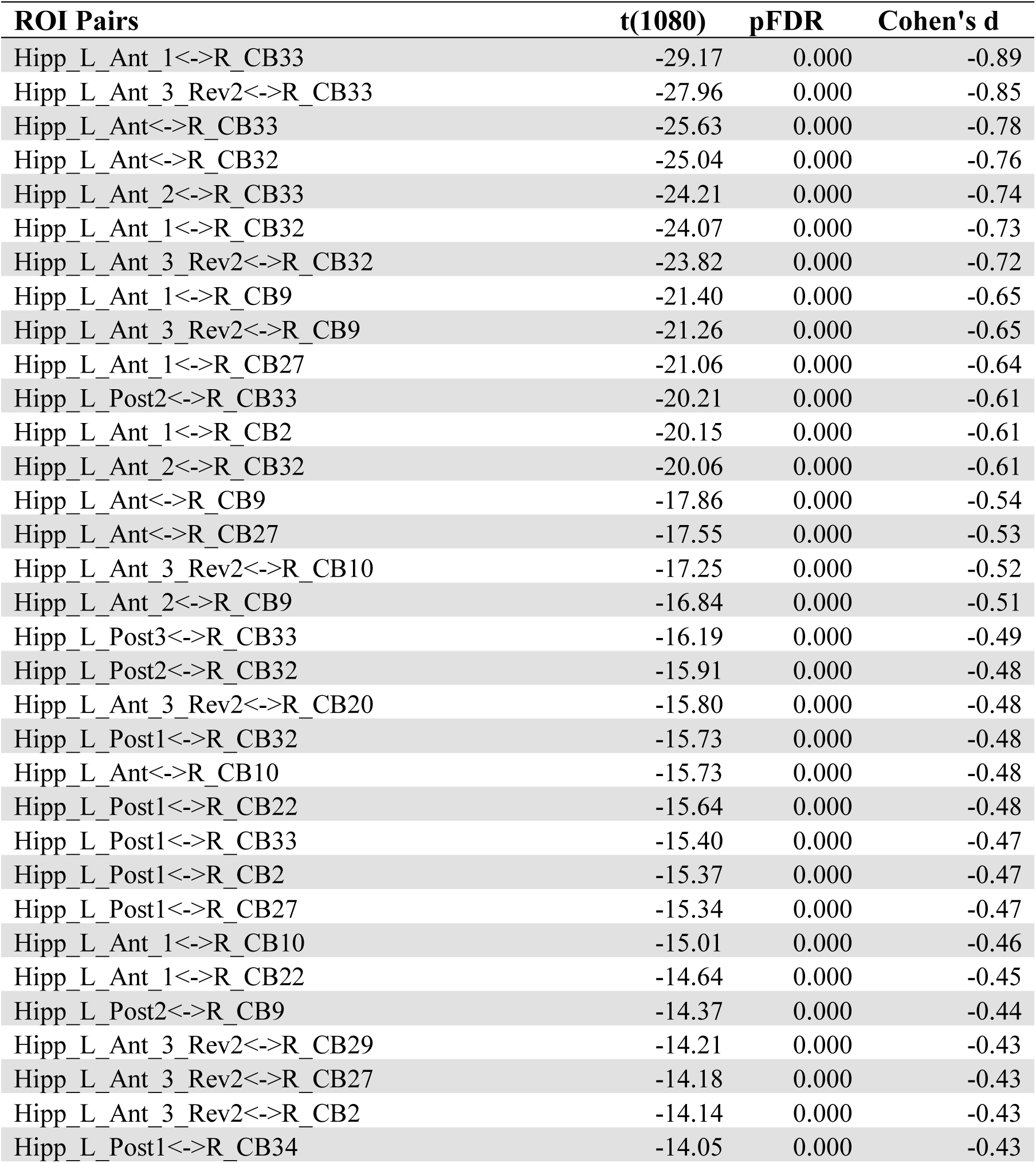
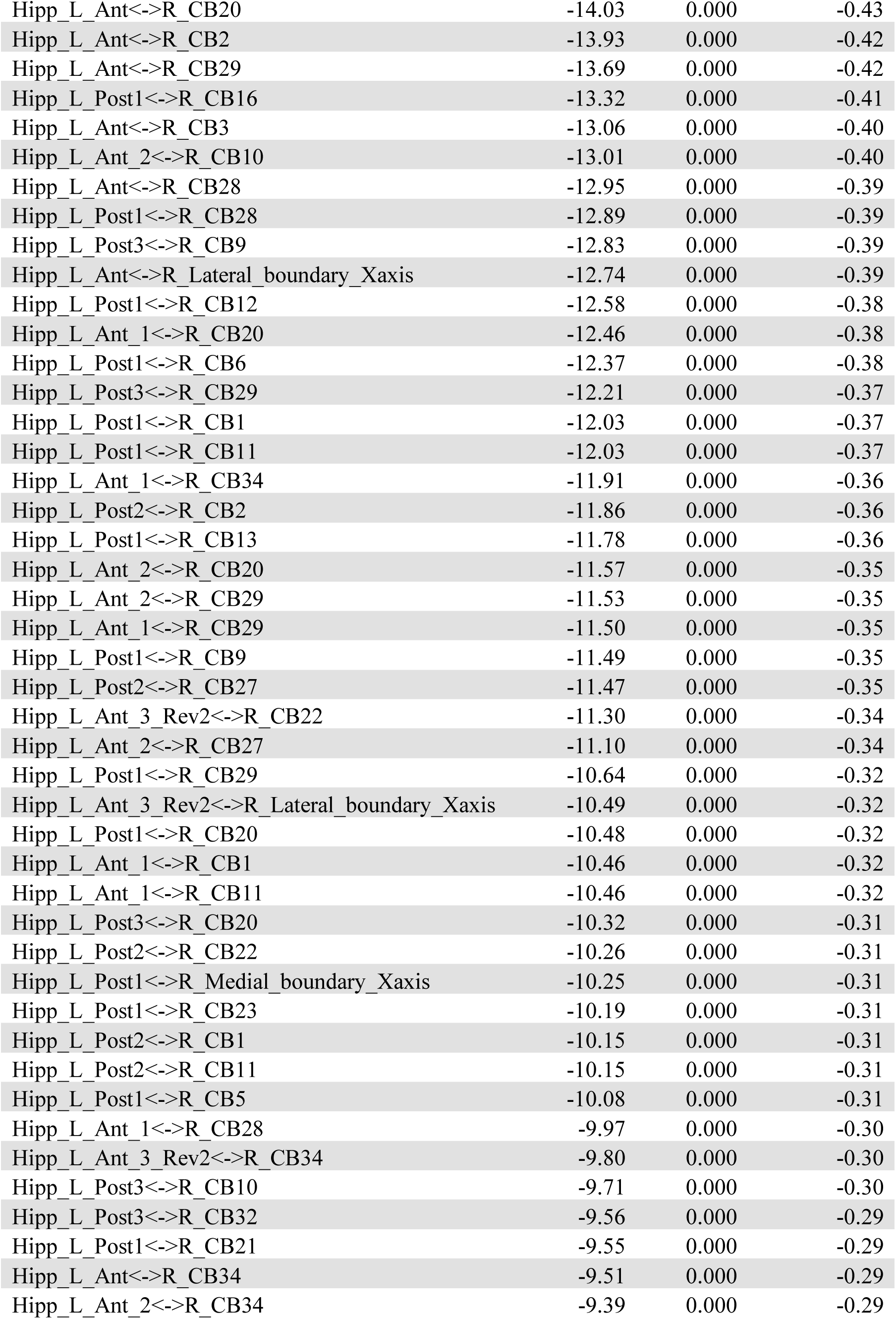
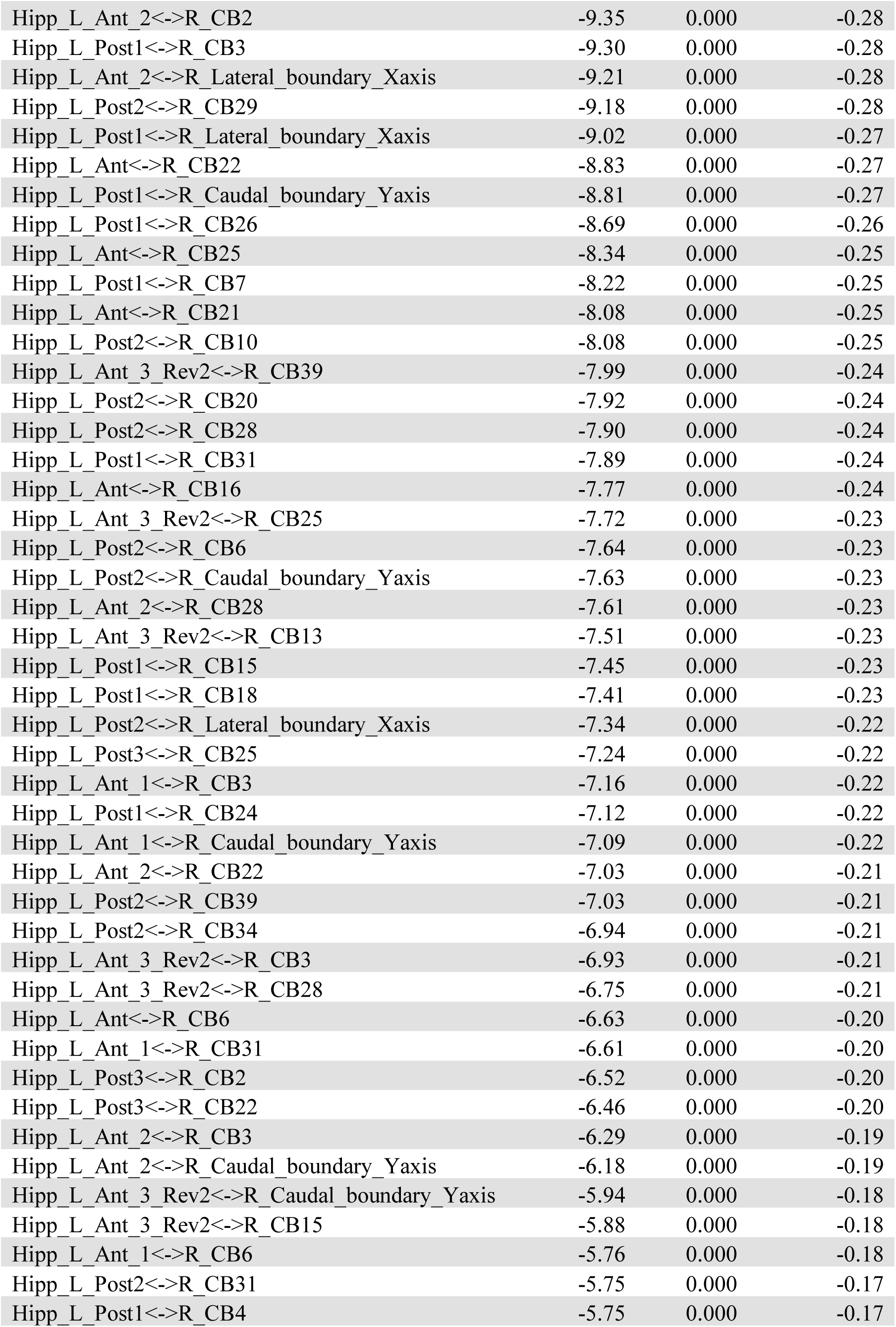
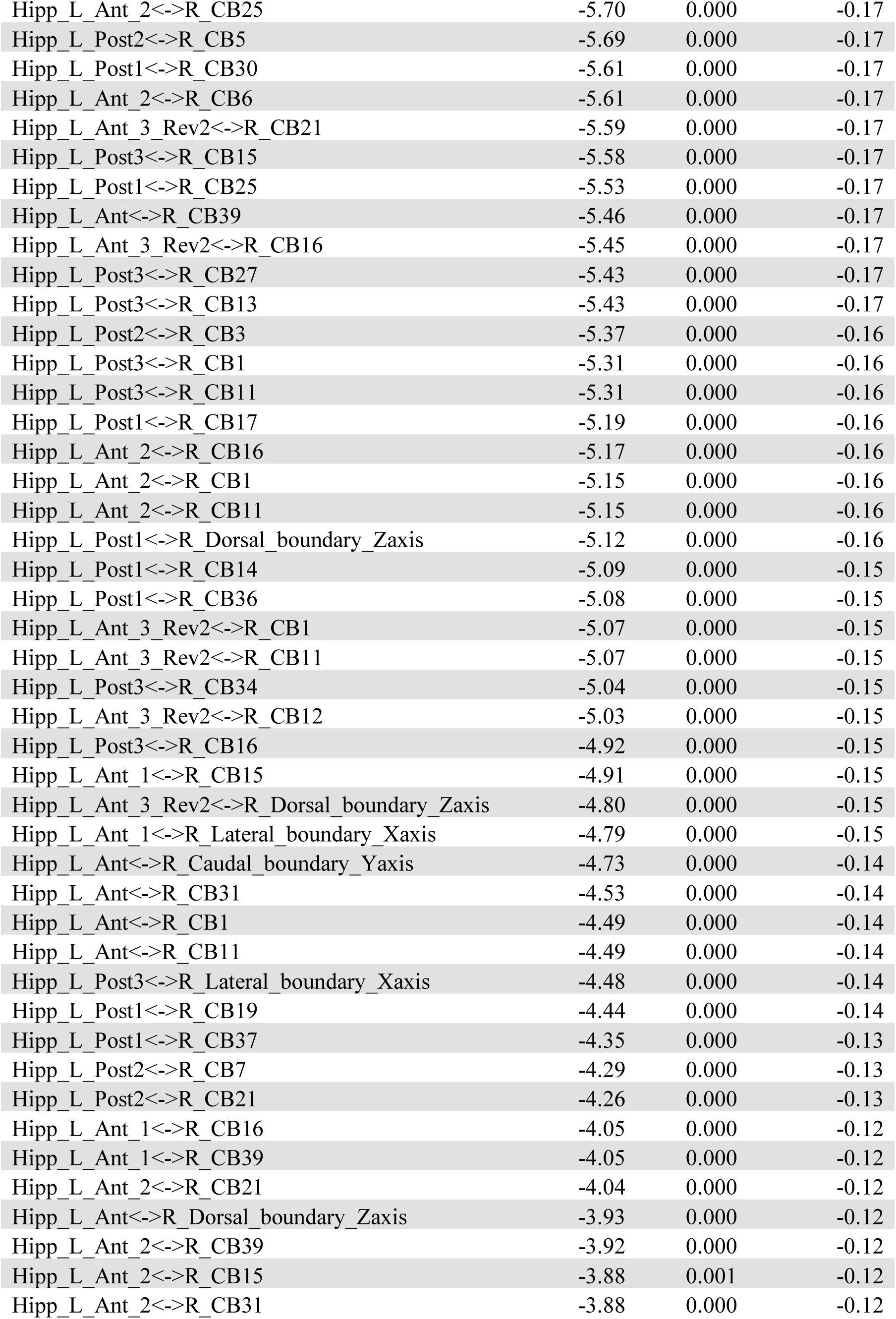
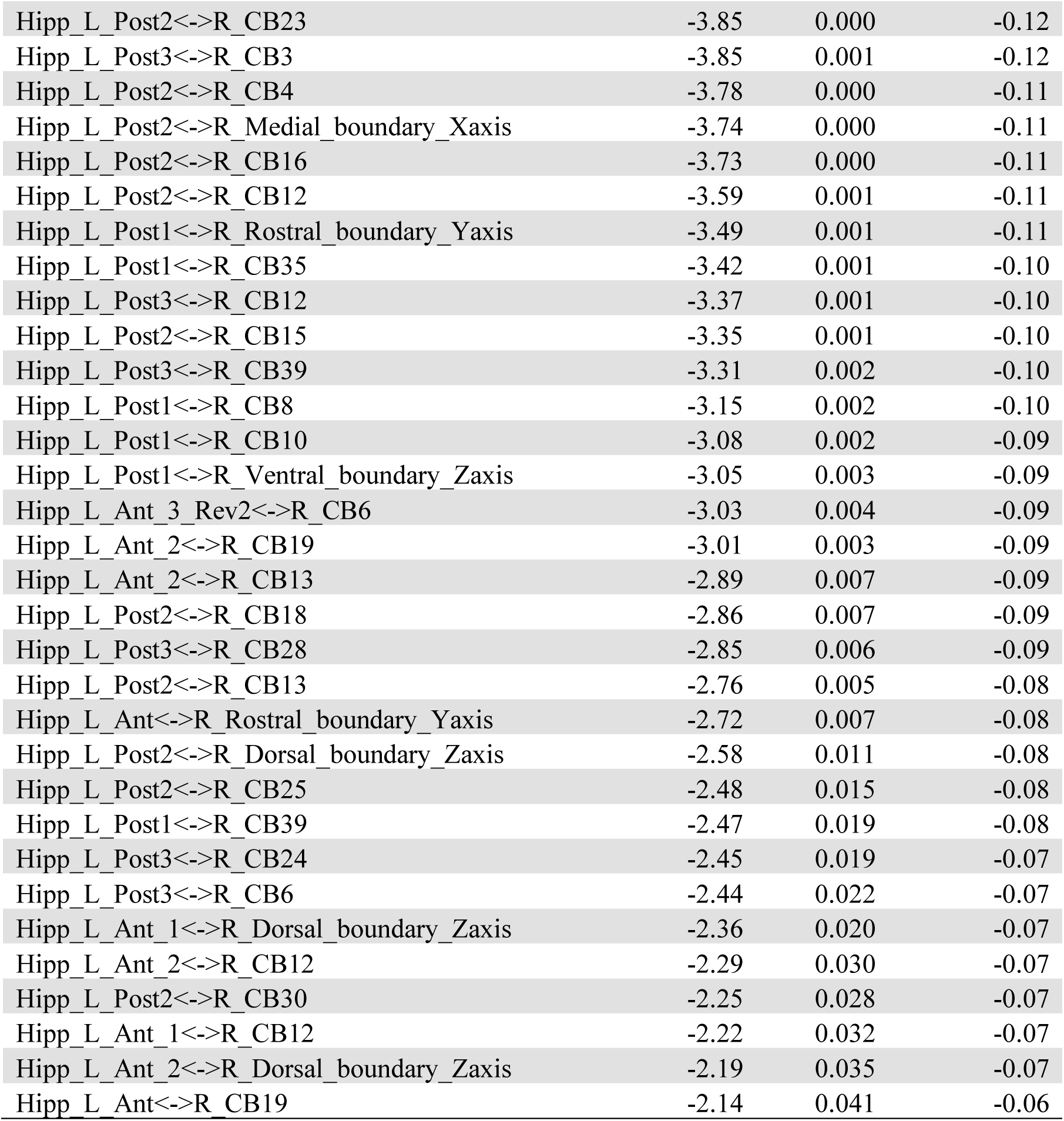
Cerebellar-hippocampal ROI pairs showing significant negative FC. In order of largest to smallest Cohen’s d effect size.

**Table 5.**
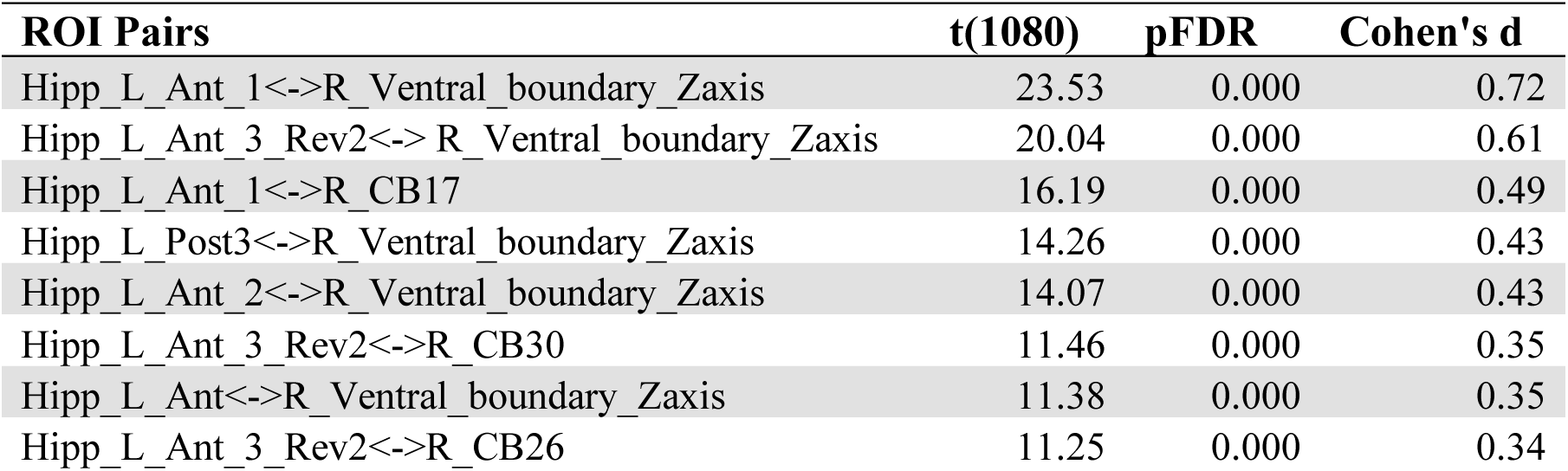
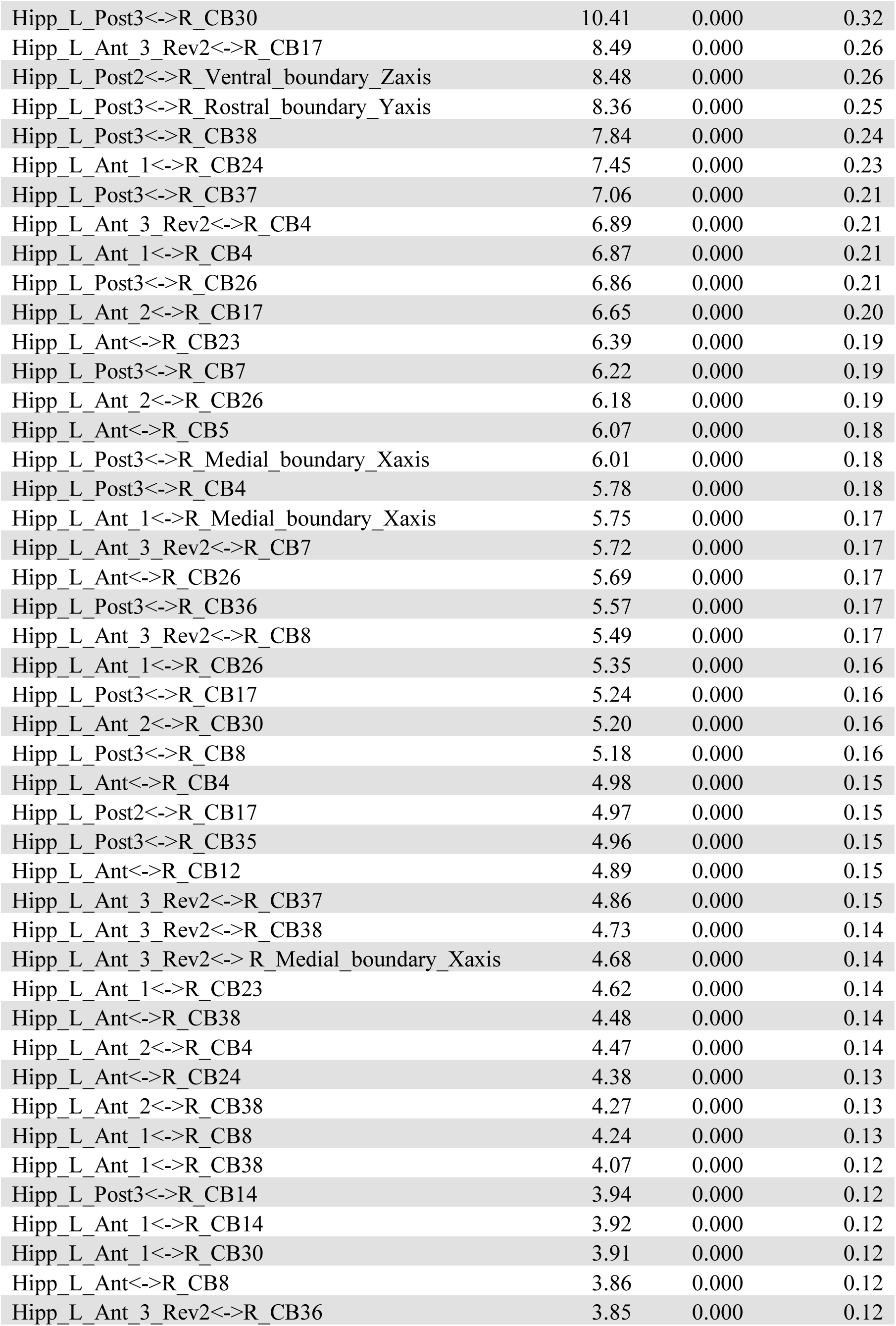
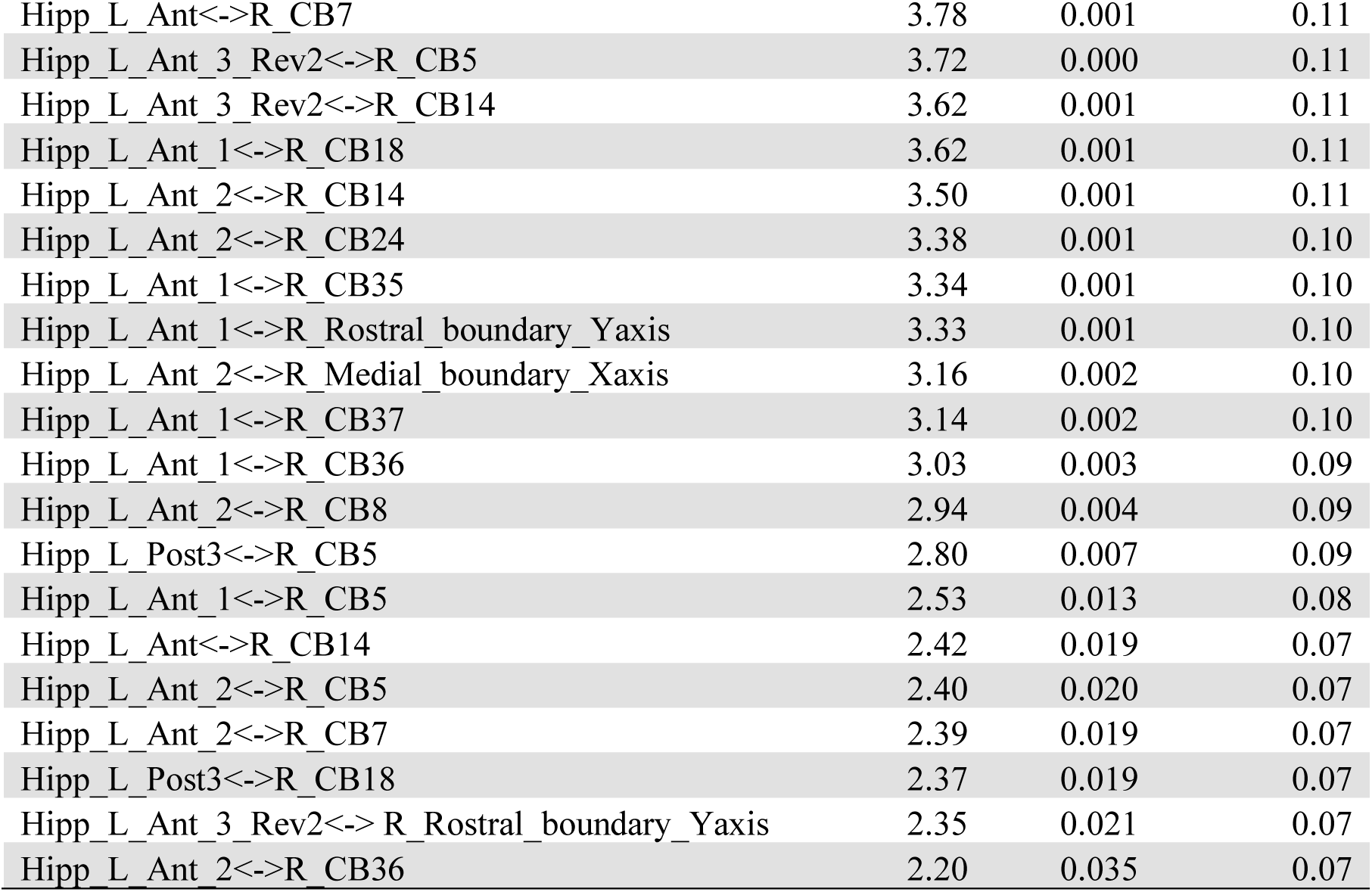
Cerebellar-hippocampal ROI pairs showing significant positive FC. In order of largest to smallest Cohen’s d effect size.

**Table 6.** Cerebellar-hippocampal ROI pairs showing significant sex difference (female > male) in CB-HP FC. In order of largest to smallest Cohen’s d effect size.

| ROI Pairs | t(1080) | pFDR | Cohen's d |
| --- | --- | --- | --- |
| Hipp_L_Ant_1<->R_Ventral_boundary_Zaxis | 5.70 | 0.013 | 0.11 |
| Hipp_L_Ant_3_Rev2<->R_CB21 | 4.39 | 0.013 | 0.08 |
| Hipp_L_Ant_3_Rev2<->R_Ventral_boundary_Zaxis | 3.83 | 0.021 | 0.07 |
| Hipp_L_Post3<->R_CB4 | 3.55 | 0.013 | 0.07 |
| Hipp_L_Ant_3_Rev2<->R_CB8 | 3.49 | 0.024 | 0.07 |
| Hipp_L_Ant<->R_Ventral_boundary_Zaxis | 3.39 | 0.013 | 0.07 |
| Hipp_L_Ant_3_Rev2<->R_CB35 | 3.24 | 0.013 | 0.06 |
| Hipp_L_Ant<->R_CB39 | 3.11 | 0.039 | 0.06 |
| Hipp_L_Ant<->R_Dorsal_boundary_Zaxis | 3.02 | 0.037 | 0.06 |
| Hipp_L_Ant_1<->R_CB31 | 3.01 | 0.034 | 0.06 |
| Hipp_L_Ant_3_Rev2<->R_Dorsal_boundary_Zaxis | 2.97 | 0.025 | 0.06 |
| Hipp_L_Post3<->R_Dorsal_boundary_Zaxis | 2.84 | 0.045 | 0.05 |
| Hipp_L_Ant_3_Rev2<->R_CB14 | 2.82 | 0.024 | 0.05 |
| Hipp_L_Ant<->R_CB4 | 2.77 | 0.025 | 0.05 |

### Cognitive Performance and CB-HP Connectivity

We investigated sex differences in cognitive performance here to contribute to extant sex difference findings^47–52^ and contextualize our CB-HP FC results. Regarding relationships across all participants between cognitive measures (i.e., Variable Short Penn Line Orientation, Penn Word Memory Test total, Penn Word Memory Test reaction time, Picture sequence memory, Flanker Task, and List Sorting Working Memory) and CB-HP FC, while controlling for age and education level, we did not find significant linear relationships (pFDR > .05). Our investigation of sex by cognitive performance interactions with CB-HP connectivity also did not yield significant sex by cognitive performance interactions as they relate to CB-HP FC revealed here (pFDR > .05).

### Sex and Cognitive Performance

Sex differences on these cognitive measures (i.e., Variable Short Penn Line Orientation, Penn Word Memory Test total, Penn Word Memory Test reaction time, Picture sequence memory, Flanker Task, and List Sorting Working Memory) have not previously been reported. We found significant sex differences for cognitive measures: Variable Short Penn Line Orientation, Penn Word Memory Test total, Picture sequence memory, Flanker Task, and List Sorting Working Memory (**Table 7**).

**Table 7.** Sex Differences in Cognitive Performance Controlling for Age and Education Level.

| Sex Differences in Cognitive performance |  |  |  |  |
| --- | --- | --- | --- | --- |
| Cognitive Measure | Sex Difference | df | F-statistic | pFDR |
| Variable Short Penn Line Orientation | M>F | 1069 | 58.82 | <.0001 |
| Picture sequence memory | F>M |  |  | 0.0001 |
| Penn Word Memory Test total | M>F | 1069 | 11.87 | 0.0009 |
| Penn Word Memory Test RT | Not Significant | 1069 | 0.05 | 0.8237 |
| Flanker task | M>F | 1076 | 16.78 | <.0001 |
| List Sorting Working Memory | M>F | 1076 | 65.63 | 0.0026 |

## Discussion

This study represents the first systematic examination of CB-HP FC in relation to age, sex, and behavioral performance in healthy young adults. Leveraging high-resolution rsfMRI and a targeted selection of cognitive tests, we examined the CB-HP circuit and tested its associations with demographic and cognitive variables. Consistent with robust evidence for direct cerebello-to-hippocampal projections ^5–7^, our resting-state analyses revealed significant CB-HP FC patterns in young adults. Further, our data showed significant sex differences in CB-HP FC. Consistent with prior reports of sex differences in cognitive performance^47–52^ and brain function^18,53^, we identified robust sex differences across the majority of our cognitive measures. These findings underscore both the specificity and limits of CB-HP functional coupling in this age range and establish a foundational map for future studies of healthy aging.

Mapping the CB-HP circuit in healthy young adults revealed a striking topography: most pairs linking the entire hippocampal axis to cerebellar territories (vermis VII, lobules VI, VII/VIIb, VIII, and Crus I/II) showed negative FC. In contrast, positive FC emerged between most hippocampal segments and ventral/medial cerebellar regions (lobules VI, VII, IX; vermis IX; Crus II). Notably, two posterior hippocampal seeds (Hipp_L_Post1, Hipp_L_Post2) were disproportionately characterized by negative or absent correlations in these same cerebellar fields, consistent with anterior-posterior gradient FC findings that have been linked to functional activation on cognitive tasks^14,54^. The rostral–ventral–lateral cerebellum encompass somatomotor and vestibular representations^54^. Widespread negative correlations of these motor-related cerebellar regions to the hippocampus as well as the contrasting positive correlations between cerebellar regions associated with ventral attention, default mode, and frontoparietal networks can be contextualized with the anterior–posterior specialization of the hippocampus^14^. That is, our findings may elucidate a motor to high-level cognition functional gradient pattern. Given proposals that cerebellar circuits scaffold cortical–hippocampal computations and may up-regulate with aging or disease (Bernard, 2024), our young-adult map defines the direction and loci of coupling prior to compensatory shifts. Collectively, the convergence of our resting topology with task findings suggests that gradients (motor to cognitive in cerebellum and gist to precision in hippocampus) organize CB–HP communication. Taken together, these patterns both support the existence of direct CB-HP pathways^5–7^ and underscore the value of a regionally resolved framework for defining a normative “hippobellum”^25^ topology in early adulthood, which is a prerequisite for detecting later hormone-, age-, or pathology-related departures from this baseline.

Females showed stronger anterior/mid-hippocampus FC with medial cerebellar regions (vermis IV–VI; lobules VI, VIII, IX; Crus II), a pattern that is consistent with reports that females generally exhibit denser local connectivity^55^, with greater default mode network (DMN) segregation as compared to males. Males show greater sensorimotor network segregation^56^ and more distributed functional organization^55^. Medial posterior cerebellar areas such as lobule IX and Crus II map onto DMN and limbic networks^54^, and hippocampal anterior/mid segments preferentially align with affective-integrative circuitry^14^, providing a network-level rationale for the female-greater FC we observed. Hormonal influences likely contribute: higher estradiol has been linked to increased large-scale network coherence and FC^21,22^. Consistent with this, our recent lifespan work shows that CB-HP FC varies with sex steroid hormone levels: greater CB-HP FC with increased estradiol and lower CB-HP FC with higher progesterone across male and female participants^12^, suggesting hormone-driven rather than sex alone differences in FC. With that said, it is critical to acknowledge that in this sample we can only speculate as hormone levels or information about menstrual cycle phase were not available.

The sex differences we observed in cognitive performance align with a growing body of evidence for sex-specific behavioral profiles. Meta-analytic work demonstrated that females display a small-to-moderate advantage in verbal episodic memory that emerges in late adolescence and remains stable through early and mid-adulthood^49^. In contrast to these findings and our hypothesis, males outperformed females on verbal episodic memory in this sample after controlling for age and education level. Voyer and colleagues found a reliable male advantage in visuospatial working memory tasks in young adults^52^ which was statistically consistent with our findings on the Variable Short Penn Line Orientation and List Sorting Working Memory tasks. Further, while males have historically outperformed females on visuospatial episodic memory^48,51^, females scored higher than males on Picture Sequence Memory in this sample. Consistent with prior work on selective attention and interference inhibition^57,58^, males in our cohort also outperformed females on the Flanker Inhibitory Control and Attention Test. Taken together, the cross-domain reversals we observed, male advantages on verbal recall and visuospatial working memory and female advantages on episodic sequence learning, suggest that sex effects are task-specific rather than uniformly favoring one group. Such heterogeneity may reflect differences in cognitive strategy, motivational salience, and/or neuroendocrine state that differentially recruit networks implicated in attention, memory, and spatial processing.

CB-HP FC was unexpectedly independent from behavior in healthy young adults. Several factors may quantify the null findings regarding CB-HP FC to behavioral associations. First, the CB-HP pathway, while anatomically confirmed via tractography^5^ and optogenetic studies^6,7^, may operate at a level of connectivity or temporal dynamics not captured by conventional resting-state analyses in a neurotypical young adult sample. Notably, FC between the cerebellum and hippocampus has been demonstrated during spatial-temporal projection of moving targets, visuospatial processing, spatial navigation, and reward-based motor learning^8–11^. Future studies should employ longitudinal designs and fMRI task-dependent paradigms to pinpoint when and how the CB-HP pathway gains behavioral relevance.

Importantly, there are several caveats to the present conclusions. First, our cross-sectional design cannot disentangle cohort effects from true intra-individual change, limiting causal inference in our CB-HP characterization as well as sex differences. Second, a mixed-handedness sample (6% left-handed and 9% ambidextrous) introduces variability because handedness modulates hand-motor, higher visual, and emotional processing^59^. While this introduces variability, it also has the advantage of being more representative of a normative population. Third, relating rsfMRI FC to offline behavioral tests may underestimate circuit relevance that only emerges under task demand; task-based fMRI stands to uncover stronger, context-specific CB-HP coupling as it has in previous CB-HP work^8–10^. Finally, our FDR correction across hundreds of ROI-to-ROI pairs reduces false positives but likely inflates false negatives, potentially obscuring subtle behavioral associations. Longitudinal, task-evoked studies with handedness-controlled samples and hypothesis-driven ROI sets will therefore be critical to validate and extend these findings.

In sum, this work delivers a detailed functional atlas of the CB-HP circuit in young adulthood and highlights sex differences in this circuit. Our findings confirm direct CB-HP pathways, a mapping which is essential for spotting hormone-, age-, or disease-related deviations later in life.

## Data Availability

Imaging and non-imaging data are available via the HCP after registering. Specifically, the following resources provide starting points to access and handle the public dataset this study is based on (”Human Connectome Project–Young Adult Study”).

All supporting data and materials are available here:

- Public data website: https://www.humanconnectome.org/study/hcp-young-adult
- Register for access: https://db.humanconnectome.org/app/template/Login.vm (simple sign-on)
- Access to restricted data (includes link to e-access application form): https://www.humanconnectome.org/study/hcp-young-adult/document/restricted-data-usage
- Data: WU-Minn HCP 1200 Subjects Data Release, for which the following reference manual applies: https://www.humanconnectome.org/storage/app/media/documentation/s1200/HCP_S1200_Release_Reference_Manual.pdf

